# Active histone modifications fine-tune DNA N-6 methyladenine deposition and maintain transcriptional stability

**DOI:** 10.64898/2026.07.21.739721

**Authors:** Carlos Lax, Macario Osorio-Concepción, Natalia Nicolás-Muñoz, Ghizlane Tahiri, Stephen J. Mondo, Vivian Ng, Eusebio Navarro, Igor V. Grigoriev, Victor Meza-Carmen, Francisco E. Nicolás, Victoriano Garre

**Affiliations:** Departamento de Genética y Microbiología, Facultad de Biología, Universidad de Murcia, 30100 Murcia, Spain; Instituto de Investigaciones Químico Biológicas, Universidad Michoacana de San Nicolás de Hidalgo, Ciudad Universitaria, 58030 Morelia, Michoacán, México; U.S. Department of Energy Joint Genome Institute, Lawrence Berkeley National Laboratory, Berkeley, CA 94720 USA; Department of Agricultural Biology, Colorado State University, Fort Collins, CO, 80523, USA; Environmental Genomics and Systems Biology Division, Lawrence Berkeley National Laboratory, Berkeley, CA, 94720, USA; Department of Plant and Microbial Biology, University of California Berkeley, Berkeley, CA 94720 USA

**Keywords:** Fungal epigenetics, histone modifications, 6mA, chromatin, *Rhizopus microsporus*

## Abstract

Epigenetic mechanisms provide sophisticated regulatory layers that modulate gene expression across diverse organisms, yet their organization and crosstalk remain poorly understood in non-dikarya fungi (NDF). Here, we characterize the genome-wide landscape of chromatin organization in the fungus *Rhizopus microsporus*, revealing a compartmentalized architecture where active histone modifications (H3K4me1, H3K4me3, H3K27ac) define transcriptionally active euchromatin distinct from H3K9me3-marked constitutive heterochromatin. Through comprehensive ChIP-seq analysis, we demonstrate that these modifications exhibit distinct distribution patterns over gene bodies and co-localize with 6-methyladenine (6mA) clusters (MACs), an essential epigenetic mark that is associated with transcription in this fungus. We identified functional specialization among H3K4 methyltransferase Set1 paralogs, where Set1a primarily deposits H3K4me3 and Set1b deposits H3K4me1. In contrast, both Gcn5 paralogs function redundantly in H3K27 acetylation. Knockout analysis reveals that these enzymes are critical for sporulation, stress resistance, and pathogenesis. Importantly, we uncover an epigenetic crosstalk in which active histone modifications restrict off-target 6mA deposition, regulate methylation cluster stability, and buffer transcriptional variation. Our findings reveal conserved principles of epigenetic crosstalk between active histone modifications and the essential DNA modification 6mA that may represent a fundamental mechanism of chromatin regulation in eukaryotes.

**SIGNIFICANCE:** Epigenetic mechanisms regulate gene activity without altering the DNA sequence, yet how different epigenetic marks interact remains poorly understood. Here, we characterize the genome-wide distribution of active histone modifications and DNA N6-methyladenine (6mA) in the fungus *Rhizopus microsporus*, revealing that they define distinct active and inactive chromatin domains. While 6mA plays a central role in transcriptional regulation, active histone modifications direct its accurate deposition and maintenance, thereby reducing transcriptional variability. These findings uncover conserved crosstalk between histone modifications and 6mA that may represent a fundamental principle of chromatin regulation across eukaryotes.

## INTRODUCTION

Epigenetic mechanisms provide a sophisticated regulatory layer that modulates gene expression without altering the underlying DNA sequence (1, 2). In eukaryotes, these mechanisms encompass diverse molecular processes, including DNA methylation, non-coding RNA-mediated regulation, chromatin remodeling, and post-translational modifications of histone proteins (3–5). Together, these systems orchestrate dynamic changes in chromatin structure and genome accessibility, enabling cells to establish and maintain distinct gene expression programs in response to developmental and environmental cues (1, 6). Importantly, epigenetic mechanisms do not function in isolation but rather engage in extensive crosstalk, establishing an epigenetic code that coordinates chromatin states and gene expression (7, 8). DNA methylation and histone modifications are intimately interconnected and mutually dependent for their establishment and maintenance (8, 9). For instance, DNA methylation can recruit histone-modifying enzymes that establish repressive chromatin marks, while certain histone modifications can direct DNA methyltransferases to specific genomic regions (7, 10). This bidirectional interplay creates self-reinforcing epigenetic states that ensure stable yet reversible gene silencing or activation (11, 12). Moreover, the interpretation of these marks by reader proteins adds a layer of complexity, as chromatin-binding domains can recognize specific combinations of DNA and histone modifications to recruit downstream effectors (13, 14).

Among histone modifications, post-translational modifications of specific lysine residues have emerged as key determinants of chromatin states and transcriptional activity (15, 16). In particular, the acetylation of histone lysine residues mediated by histone acetyltransferases (HAT) constitutes one of the main chemical marks modulating the chromatin structure, altering the interaction between DNA and histone proteins, and facilitating the access of regulatory proteins to DNA (17). The highly conserved Gcn5 (General control non-depressible-5) protein is an acetyltransferase that functions as the catalytic subunit of the SAGA and ADA coactivator complexes. Gcn5 primarily acetylates specific lysine residues on histone H3, including H3K9, H3K14, H3K27, and H3K36 (18, 19), as well as some lysine residues of the histone H4 and H2B (20, 21). It has been shown that Gcn5 is key in the control of several biological processes and virulence in human fungal pathogens. For example, in some *Candida* species, *gcn5* deletion affects lysine acetylation levels, morphogenesis, cell wall maintenance, antifungal resistance, and virulence (22). Meanwhile, in *Cryptococcus neoformans*, Gcn5 is the main player in completing the sexual cycle, sporulation, yeast-hyphal transition, hyphal development, and meiosis (23). In addition, acetylation of H3K9 and H3K14 has been associated with the transcriptional activation of master regulators involved in mating and conidiation processes (23, 24). Recent data indicate a coordinated crosstalk between histone acetylation and methylation; both systems function as critical regulators of the activation of gene transcription processes. *Candida albicans* strains lacking the methylation of histone H3 lysine 4 (H3K4) show high levels of acetylation in the same residue of histone H3, leading to gene expression in the absence of an external stimulus (25). The H3K4 methylation is performed by the histone methyltransferase Set1 (SET domain protein 1), which is associated with active transcription (26). In addition to monomethylation of H3K4 (H3K4me), Set1 protein also carries out the dimethylation and trimethylation of H3K4 (27, 28). Generally, these methylation marks are recognized by reader proteins, allowing an active state of the chromatin (13). In *Mucor lusitanicus*, a model organism for studying mucormycosis, the *set1* deletion markedly reduces H3K4 methylation, impairs sporulation, delays the filamentous growth, and attenuates virulence (29).

Fungi have been pivotal model organisms in epigenetics, contributing to fundamental discoveries in this field. The first histone deacetylase was cloned and purified in *Saccharomyces cerevisiae* (30) while studies in the fission yeast *Schizosaccharomyces pombe* revealed conserved mechanisms of H3K9 recognition and heterochromatin formation (31), as well as their inheritance mitosis and meiosis (32, 33). In *Neurospora crassa*, characterization of the histone methyltransferase Dim5 established the functional connection between histone and DNA methylation (34). Cytosine methylation in Dikarya fungi is primarily associated with transposable element control and the RIP (Repeat-Induced Point mutation) mechanism in ascomycetes and basidiomycetes (35–37). Despite these advances, the remarkable diversity of fungal epigenetic landscapes (38), together with a strong bias toward ascomycetes and basidiomycetes limits our comprehensive understanding of epigenetic regulation across the fungal kingdom. This highlights the need to investigate non-Dikarya fungi (NDF), a subkingdom encompassing the greatest number of phyla and phylogenetic diversity within fungi (39). Epigenetic mechanisms in NDF remain poorly characterized although notable contributions underscore the importance of expanding research beyond Dikarya (40). Notably, the loss of centromeric histone CENP-A in the phylum Mucoromycota represents a distinctive feature that shapes their unique centromere organization (41). Even more strikingly, NDF commonly employ 6-methyladenine (6mA) as the primary DNA modification in contrast to the predominance of 5-methylcytosine in most eukaryotes (42–45). In several NDF representatives, 6mA occurs at high levels (up to 2-3% of total adenines) and predominantly as a symmetric modification at ApT dinucleotides. In contrast, 6mA is nearly absent in most eukaryotes, where the low levels of methylation that are detected occur predominantly in an asymmetric context (42, 44, 46, 47). Its enrichment in actively transcribed genes and its depletion from poorly expressed or silenced genes suggest a role in transcriptional regulation. Together with its essential function in the mucoralean fungus *Rhizopus microsporus* and its pervasive involvement in the regulation of gene expression (43, 48), Symmetric 6mA emerges as a central component of the epigenetic landscape of this fungus. Interestingly, 6mA distribution is associated with a compartmentalized chromatin organization, where it colocalizes with regions marked by H3K4me3 and is excluded from constitutive heterochromatin domains enriched in H3K9me3 (43).

Here, we further explore the compartmentalized genome architecture of *R. microsporus*, uncovering the distribution and functional roles of major euchromatin histone modifications and the enzymatic machinery responsible for their deposition. We show that H3K4me1, H3K4me3, and H3K27ac delineate distinct euchromatic subdomains coordinated by Set1 and Gcn5 paralog complexes. Notably, the loss of these active histone marks does not globally disrupt 6mA but instead redistributes it to ectopic genomic regions, revealing a role for active histone modifications in constraining the genomic landscape of DNA 6mA.

## RESULTS

### Active Chromatin Marks Define Open Chromatin Sub-Compartments in *R. microsporus*

To characterize the genome-wide landscape of chromatin organization at the epigenetic level in *R. microsporus* and its interplay with gene expression regulation, we generated ChIP-seq data for histone methylation (H3K4me1 and H3K4me3), acetylation (H3K27ac), and RNA polymerase II (RNA Pol II). The three modifications showed a similar global distribution, appearing in gene-rich and transcriptionally active regions and absent in repeat-rich H3K9me3 constitutive heterochromatin regions (**Figure 1A and Supplementary Figure 1A)**. Consequently, *R. microsporus* transposable elements (TEs) appeared devoid of these modifications, while they showed distinctive distribution patterns over genes (**Figure 1B**). H3K4me3 preferentially accumulated downstream of the transcription start site (TSS), coinciding with previous observations (43), whereas H3K4me1 and H3K27ac preferentially accumulated upstream of the TSS, decayed right at the TSS, and increased along the gene body up to the transcription termination site (TTS) (**Figure 1B**). Genome-wide comparison of their distribution revealed two major chromatin compartments (**Figure 1C**): a constitutive heterochromatin compartment enriched in H3K9me3 and predominantly associated with TEs and lncRNAs (43, 49), and an open chromatin compartment that could be further subdivided into H3K4me3/6mA-enriched and H3K4me1/H3K27ac-enriched regions. Furthermore, H3K4me3 largely co-localized with MACs (Methylated Adenine Clusters), whereas H3K4me1 and H3K27ac did not (**Figure 1D**), reflecting their distinct distribution across gene bodies and flanking regions (**Figure 1B**).

**Figure 1.**
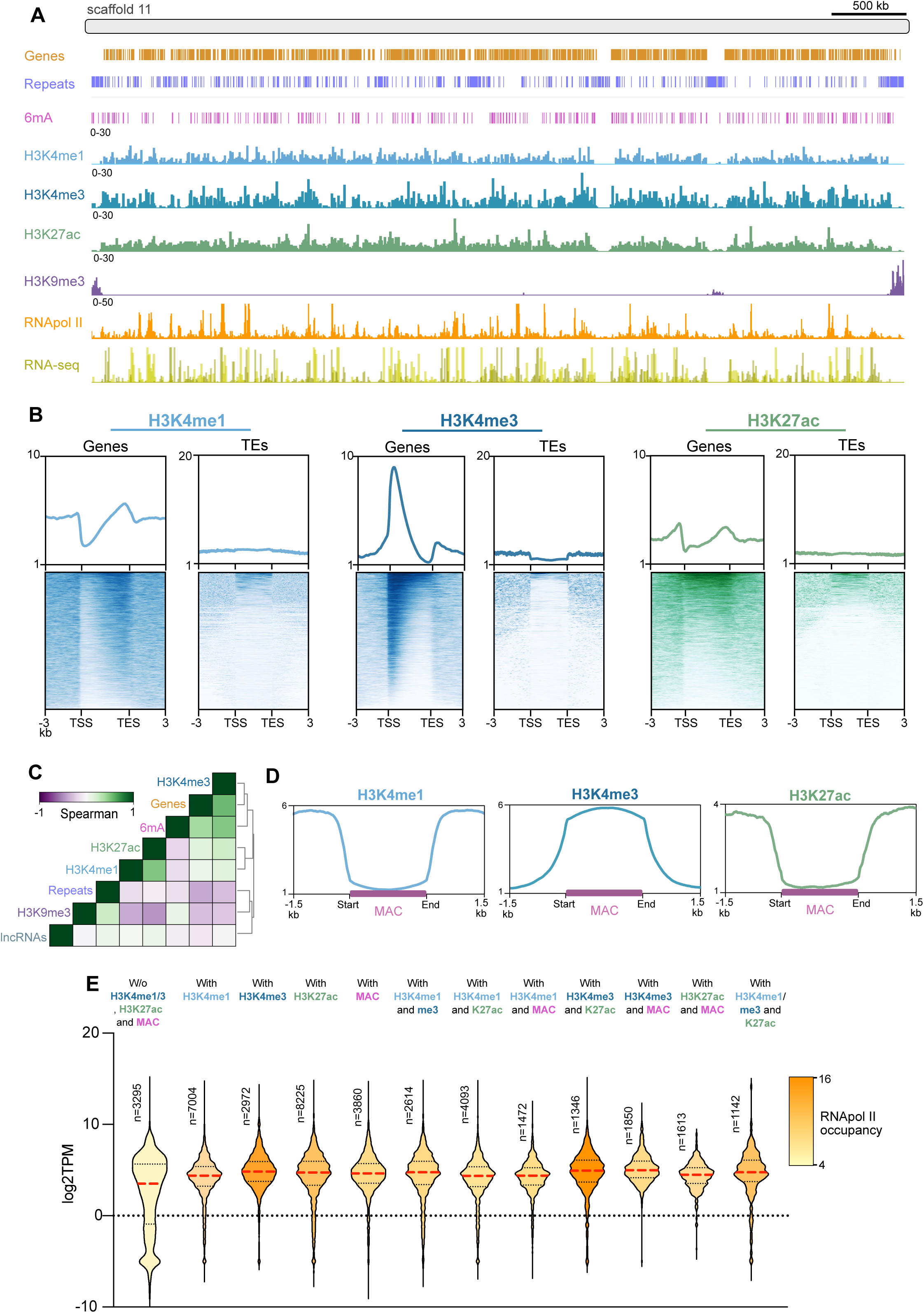
Genome-wide landscape of chromatin organization and its association with transcriptional activity in *R. microsporus*. (**A**) Genomic tracks of Genes (light brown), Repeats (purple-blue), 6mA (pink), H3K4me1 (light blue), H3K4me3 (dark blue), H3K27ac (green), H3K9me3 (dark purple), RNA Pol II (orange), and RNA-seq (yellow) across scaffold 11. Scale = 500 kb. (**B**) Profile metaplot and heatmaps for genes and TEs showing H3K4me1, H3K4me3, and H3K27ac occupancy. Genes were sized to 4 kb and occupancy was also computed 3 kb upstream and downstream the gene body (**C**) Spearman’s correlation coefficient of genome-wide distribution of DNA and histone epigenetic features. (**D**) H3K4me1, H3K4me3 and H3K27ac occupancy over MACs. Each MAC was extended to 2 kb and fragmented into 200 bins (n = 7441). (**E**) Correlation between chromatin marks and gene expression. Genes were grouped based on the presence of H3K4me1, H3K4me3, H3K27ac, MACs, or combinations thereof. Inner Violin colors show RNA Pol II occupancy for each group.

To correlate the distribution of these modifications with regulation of gene expression, we divided the genes into four quartiles according to their expression level. As expected, RNA Pol II occupancy positively correlated with transcriptional activity, showing strong enrichment from the TSS to the TTS in highly expressed genes, whereas occupancy was markedly reduced in the quartile of poorly expressed and silent genes (Wilcoxon rank, *p* < 0.001, **Supplementary Figure 1B**). Using the same expression quartiles, we found that H3K4me1, H3K4me3, and H3K27ac were preferentially enriched at highly expressed genes and were largely absent from transcriptionally inactive genes displaying low RNA Pol II occupancy (Wilcoxon rank, *p* < 0.001, **Supplementary Figure 1C**).

To further investigate the relationship between chromatin features and transcription, we identified peaks for each epigenetic mark and classified *R. microsporus* genes according to the presence of H3K4me1, H3K27ac, MACs, and any combination of these marks. Genes lacking all these transcription activation-associated modifications exhibited significantly lower expression levels than marked genes (Kolmogorov-Smirnov Test *p* < 0.001, **Figure 1E**), although no evidence of synergistic effects on gene expression was observed among the different combinations of marks analyzed (**Figure 1E**). Consistent with these findings, genes marked with H3K4me1, H3K4me3, H3K27ac, or MAC displayed significantly higher RNA Pol II occupancy than unmarked genes, with the strongest association observed for genes carrying H3K4me3 peaks downstream of the TSS (Kolmogorov-Smirnov Test, *p* < 0.001, **Figure 1E**). Taken together, these results define the epigenetic landscape of open chromatin regions in *R. microsporus*, highlighting the distinct genomic distribution of activating histone modifications and their relationship with 6mA and transcriptional activity.

### Differential Roles of Set1 and Gcn5 Paralogs in Histone Methylation and Acetylation

To characterize in detail the biological implications of these epigenetic modifications, we identified the components responsible for their deposition in *R. microsporus*. Using the well-characterized *S. cerevisiae* Set1 protein as a query sequence, we identified two putative Set1 orthologs (Set1a, ID: 1843915; Set1b, ID: 1796522) in this fungus (**Figure 2A**), in contrast to the closely related species *M. lusitanicus*, which possesses a single Set1 ortholog (29). One of the *R. microsporus* Set1-like proteins, Set1a, contains a large Na+/Ca+ exchange domain and an RNA recognition motif in addition to the conserved SET domain (**Figure 2A**). To determine the contribution of these proteins to H3K4 methylation, we generated knockout mutants using CRISPR/Cas9 (**Supplementary Figures 2A and 2B**) and analyzed global methylation levels by western blots. In *M. lusitanicus*, the single Set1, which is phylogenetically related to *R. microsporus* Set1a, is responsible for H3K4 mono-, di-, and trimethylation. Strikingly, disruption of *set1a* resulted in a clear reduction of H3K4me3, but not a significant reduction in global H3K4me1 and H3K4me2 levels, indicating that Set1a predominantly contributes to H3K4 trimethylation (**Figure 2B**). In contrast, *set1b* disruption reduced only global H3K4me1 levels, but did not significantly reduce global H3K4me2 and H3K4me3 levels (**Figure 2B**), indicating that Set1b is the primary contributor to global H3K4 monomethylation in *R. microsporus*. The fact that neither of the two mutants showed reductions in global H3K4me2 levels suggests the involvement of additional methyltransferases or regulatory factors in the deposition of this modification. In addition, these results reveal a functional specialization of the two Set1 paralogs, representing a clear divergence from the single multifunctional Set1 enzyme described in Mucorales and most other fungi.

**Figure 2.**
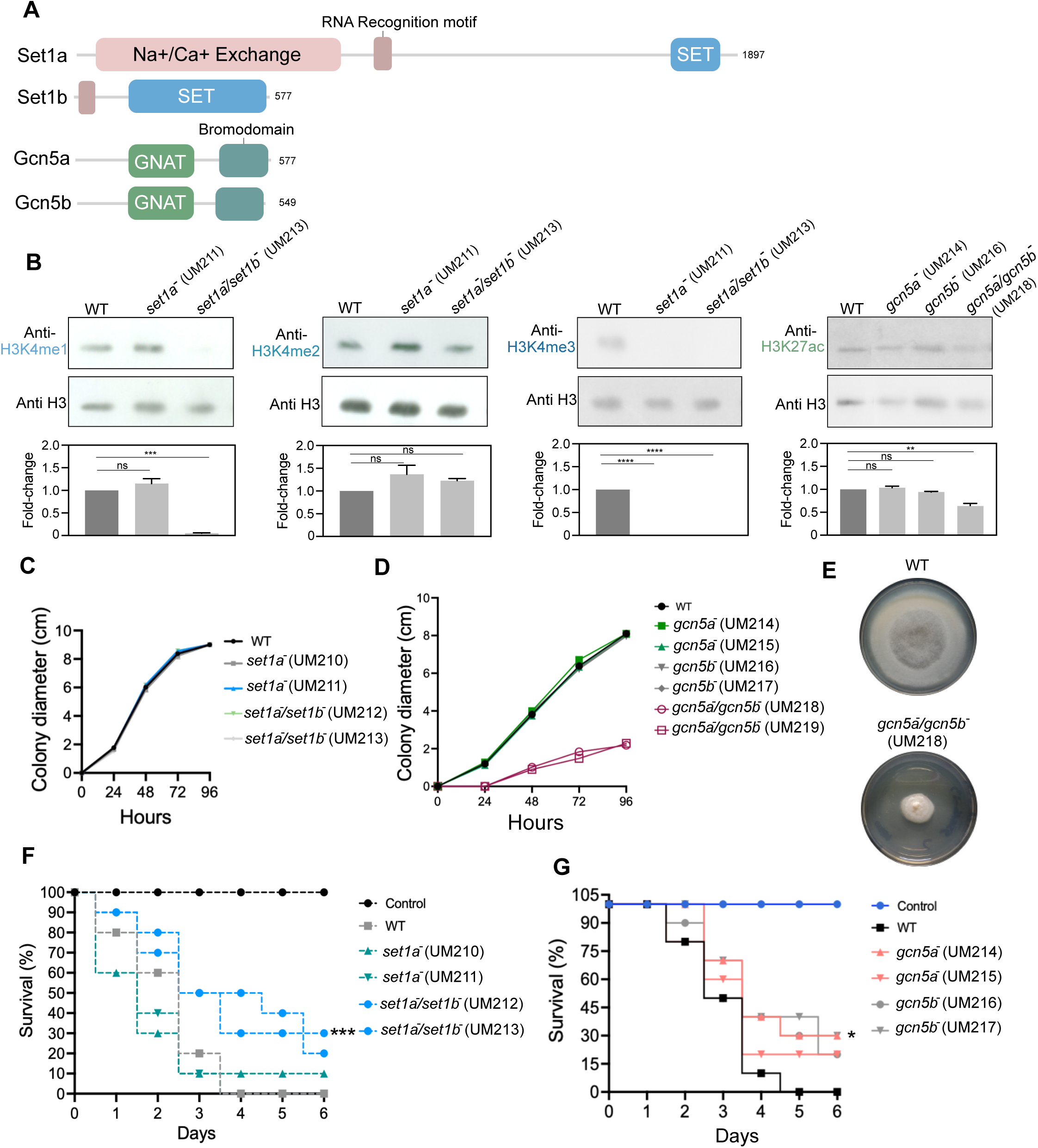
Identification and characterization of histone modification machinery. **(A)** Schematic representation of the protein sequences of Set1 and Gcn5 proteins identified in *R. microsporus*. Annotated domains are indicated, along with the total number of amino acids in each protein. (**B**) Relative quantification by Western blot using specific antibodies against H3K4me1, H3K4me2, H3K4me3, and H3K27ac. The anti-H3 antibody was used as a loading control and to normalize the signal, using the WT strain as the reference for relative densitometric comparison performed with ImageJ. Three independent replicates (from different experiments and membranes) were analyzed, and data are represented as mean ± SD. Welch’s t-test: H3K4me1 (WT vs. *set1a*^-^/*set1b*^-^, *p* = 0.0007); H3K4me3 (WT vs. *set1a*^-^ and *set1a*^-^/*set1b*^-^, *p* < 0.0001); H3K27ac (WT vs. *gcn5a*^-^/*gcn5b*^-^, *p* = 0.003). (**C**) Radial growth of *set1* mutants measured at 24, 48, 72, and 96 hours post-inoculation in YPG medium. (**D**) Radial growth of *gcn5* mutants measured at 24, 48, 72, and 96 hours post-inoculation. (**E**) Representative image of WT and *gcn5a*^-^ /*gcn5b*^-^ (UM218) in YPG medium after 72 hours post-inoculation. Defective growth and absence of aerial mycelium development can be observed in the mutant (**F**) and (**G**) survival curves for *G. mellonella* larvae inoculated with Set1 and Gcn5 mutants, respectively. The survival rate was compared with the IPS-injected larvae using the Mantel-Cox test (UM212 *p* < 0.01, UM217 *p* = 0.04)

We next investigated the Gcn5 family. Two Gcn5 possible paralogs (Gcn5a ID: 1877669 and Gcn5b ID:1894499) were identified using the well-characterized *S. cerevisiae* Gcn5 protein as a query in a BLAST search. Disruption of either *gcn5a* or *gcn5b* alone had no significant effect on global H3K27ac levels, whereas simultaneous deletion of both genes caused a pronounced reduction (**Figure 2B**). These results indicate that, unlike the functional specialization observed for the Set1 paralogs, Gcn5a and Gcn5b perform largely redundant functions in H3K27 acetylation.

### Histone Modification Machinery Regulates Critical Processes in *R. microsporus* Biology

To unravel the biological implications of the *set1* and *gcn5* genes, we conducted a comprehensive phenotypic characterization of the generated mutants. Null single and double mutants for *set1a* and *set1b*, as well as the null single mutants for *gcn5a* and *gcn5b*, showed no differences in growth, assessed by colony diameter, compared to the wild-type strain (**Figures 2C and 2D**). In contrast, the *gcn5a*^−^*/gcn5b*^−^ double mutant exhibited severely impaired growth, altered colony morphology, and a complete loss of sporangiophore production (**Figures 2D and 2E**), precluding further phenotypic analyses. Exposure to cell wall and membrane stress revealed increased sensitivity to SDS in the *set1a*^−^, *set1a*^−^*/set1b*^−^, and *gcn5b*^−^ mutants (**Supplementary Figures 2C, 2D, 2F, and 2G**). Conversely, the *set1a*^−^*, gcn5a*^−^, and *gcn5b*^−^ mutants displayed enhanced resistance to oxidative stress (**Supplementary Figures 2C, 2E, 2F, and 2H**). Finally, virulence assays using *Galleria mellonella* larvae showed that the *set1*^−^ double mutants and *gcn5b*^−^ single mutant were significantly less virulent than the wild-type strain (**Figures 2F and 2G**)

### Loss of Active Histone Marks Correlates with Reduced RNA Pol II Occupancy and Gene Downregulation

To investigate how loss of histone methylation and acetylation affects transcriptional regulation in *R. microsporus*, we analyzed genome-wide changes in chromatin modifications and gene expression in the mutant strains. ChIP-seq confirmed the loss of H3K4me1 and H3K4me3 in the *set1a*^−^/*set1b*^−^ double mutant and a partial reduction in H3K27ac across the genome in the *gcn5a*^−^/*gcn5b*^−^ double mutant (**Figure 3A and Supplementary Figure 3A**). We next examined the transcriptional consequences of losing these activating chromatin marks. Genes that lost H3K4me1, H3K4me3, or H3K27ac peaks showed a strong bias towards downregulation with 85.2%, 91.2%, and 75.7% of differentially expressed genes (DEGs), respectively, exhibiting reduced expression (**Figure 3B**), further supporting the association of these epigenetic modifications with transcriptional activation. Consistent with the extensive co-localization of H3K4me1, H3K4me3, and H3K27ac within open chromatin regions, the *set1a*^−^*/set1b*^−^ and *gcn5a*^−^*/gcn5b*^−^ double mutants shared a large proportion of DEGs (**Figure 3C**). Most of these common targets were downregulated in both mutants, accounting for 71.1% and 50.4% of all DEGs in the *set1a*^−^/*set1b*^−^ and *gcn5a*^−^*/gcn5b*^−^ double mutants, respectively (**Figure 3D**), indicating that H3K4me1, H3K4me3, and H3K27ac cooperate to maintain the expression of a common set of genes.

**Figure 3.**
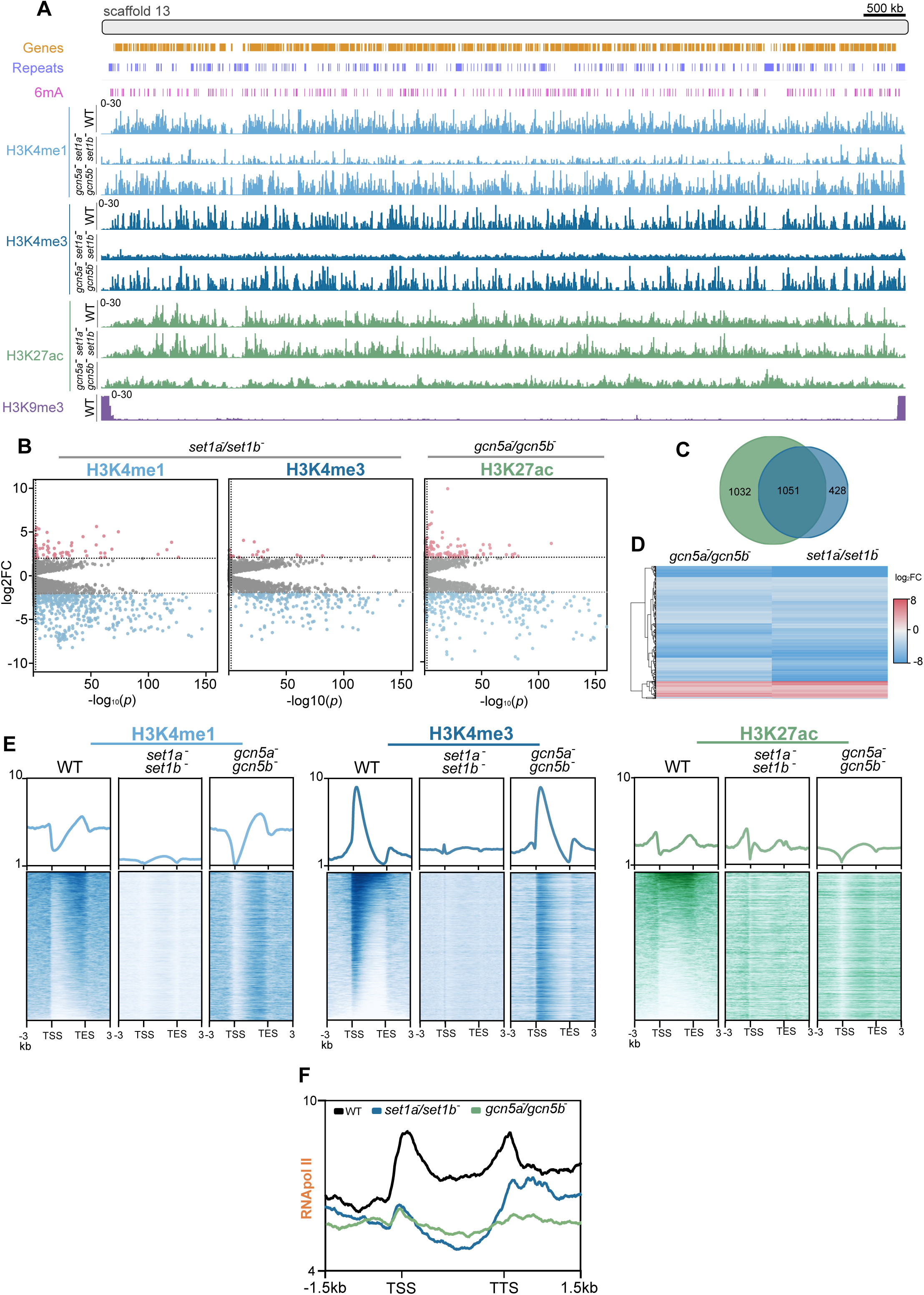
Dynamics of activating histone modification alter gene expression. **(A)** Genomic tracks of Genes (light brown), Repeats (purple-blue), 6mA (pink), H3K4me1 (light blue), H3K4me3 (dark blue), H3K27ac (green), and H3K9me3 (dark purple) for wild-type (WT) strain and double *set1* and *gcn5* mutants across scaffold 13. Scale = 500 kb (**B**) Volcano plots showing upregulated (red) and downregulated (Blue) genes that have lost an H3K4me1 or H3K4me3 peak in the *set1a*^-^/*set1b*^-^ mutant or an H3K27ac peak in the *gcn5*a^-^/*gcn5b*^-^ mutant. Only genes with FDR < 0.05 and absolute log_2_ fold change (|log₂FC|) ≥ 2 were considered as differentially expressed. (**C**) Venn diagram of DEGs in the *gcn5*a^-^/*gcn5b*^-^ mutant (green circle) and the *set1a*^-^/*set1b*^-^ mutant (blue circle). The number of DEGs is indicated on each circle and in the join zone. (**D**) Heatmap showing the log_2_FC of each common DEG between *gcn5*a^-^/*gcn5b*^-^ and *set1a*^-^/*set1b*^-^ mutants compared to the wild-type strain. (**E**) Gene profile plots and heatmaps of H3K4me1, H3K4me3, and H3K27ac occupancy in wild-type and *gcn5*a^-^/*gcn5b*^-^ and *set1a*^-^/*set1b*^-^mutants. Genes were sorted by decreasing average occupancy of H3K4me3 in the wild-type strain. Genes were sized to 4 kb and occupancy was also computed 3 kb upstream and downstream of the gene body. (**F**) RNA Pol II occupancy over the 1051 commonly DEGs between *gcn5*a^-^/*gcn5b*^-^ and *set1a*^-^/*set1b*^-^ mutants compared to the wild-type strain. Profiles are shown for WT (black), *set1a*^-^/*set1b*^-^ mutant (blue), and *gcn5*a^-^/*gcn5b^-^* mutant (green). Genes were sized to 2 kb and occupancy was also computed 1.5 kb upstream and downstream of the gene body.

Gene-wide metagene analysis confirmed the depletion of H3K4me1 and H3K4me3 across genes in the *set1a*^−^*/set1b*^−^ mutant as well as a marked reduction of H3K27ac in the *gcn5a*^−^*/gcn5b*^−^ mutant, particularly at the promoter region upstream of the TSS (**Figure 3E**). Interestingly, loss of Gcn5 function in the *gcn5a*^−^*/gcn5b*^−^ mutant also resulted in reduced H3K4me1 occupancy around the TSS, suggesting crosstalk between H3K27ac and H3K4 methylation (**Figure 3E**). The concomitant reduction of H3K27ac and H3K4me1 at promoter regions likely contributes to the widespread transcriptional repression and severe growth defects observed in the *gcn5a^-^/gcn5b^-^* mutant.

To determine whether these transcriptional changes were associated with altered recruitment of the transcriptional machinery, we conducted RNA Pol II ChIP-seq. Compared to the wild-type strain and the *set1a*^−^*/set1b*^−^ mutant, the *gcn5a*^−^*/gcn5b*^−^ mutant exhibited a genome-wide reduction in RNA Pol II occupancy across protein-coding genes (Kolmogorov-Smirnov test, *p* < 0.001; **Supplementary Figure 3B**). Likewise, the set of genes commonly differentially expressed in both mutants displayed lower RNA Pol II occupancy than unaffected genes (Kolmogorov-Smirnov test, *p* < 0.001; **Figure 3F**). Together, these findings show that disruption of activating histone methylation and acetylation impairs RNA Pol II recruitment, leading to widespread transcriptional repression.

### Histone Modifications Restrict Off-Target 6mA and Stabilize Epigenetic and Transcriptional States

To obtain a comprehensive overview of the epigenetic landscape and investigate potential crosstalk between histone modifications and DNA methylation, we conducted 6mA profiling by PacBio sequencing of the wild-type strain and the *set1a*^−^/*set1b*^−^ and *gcn5a*^−^/*gcn5b*^−^ mutants. Global genomic 6mA levels remained largely unchanged in both mutants (**Figure 4A**), reinforcing the notion that 6mA constitutes a stable epigenetic modification (50). However, the *set1a*^−^/*set1b*^−^ mutant exhibited a reduced number of MACs (**Figure 4B**), suggesting a role of H3K4 methylation in the maintenance of MACs. Global levels of symmetric 6mA (fully methylated sites) also remained largely unchanged, with only slight reductions in both mutants (83.5% in the wild-type strain versus 81.8% and 81.4% in the mutants, respectively; **Figure 4C**).

**Figure 4.**
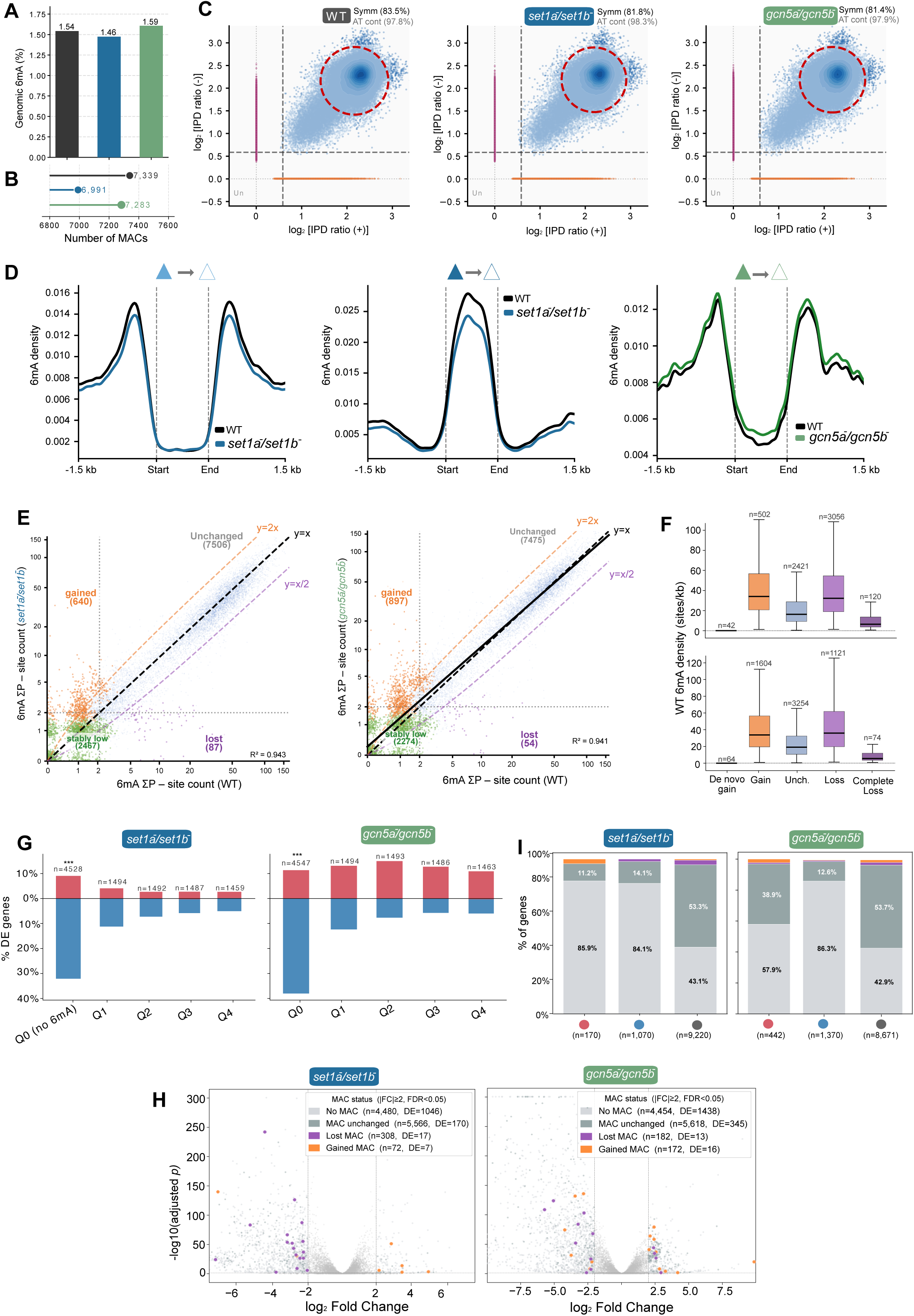
Crosstalk between histone modifications and 6mA in *R. microsporus*. (**A**) Global 6mA density expressed as the percentage of methylated adenines over total genomic adenines in WT, *set1a*⁻/*set1b*⁻ and, *gcn5a*⁻/*gcn5b*⁻ strains. (**B**) Total number of MACs detected in each strain. (**C**) Two-dimensional scatter plots of the log_2_-transformed IPD ratio for all fully symmetric 6mA sites (simultaneously methylated on both strands at A/T dinucleotides, blue) and hemimethylated (only one strand methylated, orange or purple) in each strain. (**D**) 6mA density profiles over genes that lost H3K4me1, H3K4me3, and H3K27ac peaks (filled triangle → empty triangle) in the corresponding mutant relative to WT. Wilcoxon signed-rank test *p* < 0.001 for H3K4me3. **(E**) Scatter plots of 6mA density (methylated sites per kb) in WT strain versus *set1a*⁻/*set1b*⁻ (left) and *gcn5a*⁻/*gcn5*b⁻ (right). Genes are colored by change category: Gained (orange), lost (pink), unchanged (grey), and stably low (light grey). (**F**) Boxplots of WT 6mA density for each change category in *set1a*⁻/*set1b*⁻ (top) and *gcn5a*⁻/*gcn5b*⁻ (bottom) mutants. (**G**) Percentage of upregulated (red) and downregulated (blue) DEGs (|log₂FC| ≥ 2, FDR < 0.05) stratified by 6mA density quartile for *set1a*⁻/*set1b*⁻ (left) and *gcn5a*⁻/*gcn5b*⁻ (right). Fisher’s exact test no MAC vs MAC genes *p* = 9.9 × 10^-300^ for *set1a*⁻/*set1b*⁻ and *p* = 1.6 × 10^-205^. (**H**) Volcano plots of gene expression in the *set1a*⁻/*set1b*⁻ and *gcn5a*⁻/*gcn5b*⁻ mutants. Genes are colored by their MAC status in the corresponding mutant versus the wild-type strain: No MAC (light grey), MAC unchanged (dark grey), MAC loss (orange), and MAC gain (red). Dashed lines indicate significance thresholds (|log₂FC| = 2; FDR = 0.05). The number of genes per category and the number of DEGs per category are indicated in the legend. (**I**) Proportion of genes exhibiting MAC gain (yellow), MAC loss (purple), unchanged MAC status (grey), or no MAC (light grey) among upregulated (red dot), downregulated (blue dot) and not DE genes (grey).

Given our previous findings that loss of symmetric 6mA is associated with reduced H3K4me3 occupancy over genes (43), we assessed whether the converse relationship also holds. Gene regions that lost H3K4me1 peaks in the *set1a*^−^/*set1b*^−^ mutant or H3K27ac peaks in the *gcn5a*^−^/*gcn5b*^−^ mutant showed minimal changes in 6mA distribution, whereas gene regions depleted of H3K4me3 peaks in the *set1a*^−^/*set1b*^−^ mutant exhibited a modest but significant reduction in 6mA levels (Wilcoxon < 0.01; **Figure 4D**). Therefore, most genes remained stable with no significant changes in 6mA (stably low: low modification density with no significant changes) in both mutants. However, both mutants exhibited a clear tendency to gain ectopic 6mA sites (**Figure 4E**), consistent with previous observations in ciliates showing that H3K4me3 restricts off-target 6mA deposition (50). Our results further extended this concept by suggesting that H3K4me1 and H3K27ac also contribute to maintaining the fidelity of 6mA localization. Interestingly, shorter genes with fewer methylated sites were more susceptible to alterations in 6mA distribution and complete loss of 6mA was typically observed in less densely methylated genes, whereas densely methylated genes exhibited greater stability (**Figure 4F and Supplementary Figure 4A**). We next examined adenine methylation changes at the level of entire MACs. In addition to the expected depletion in H3K4me1 and H3K4me3 levels in the *set1a*^−^/*set1b*^−^ mutant, we also found that MACs lost in the *gcn5a*^−^/*gcn5b*^−^ double mutant displayed higher-than-average H3K27ac occupancy in the wild-type strain (**Supplementary Figures 4B and 4C**), suggesting that H3K27ac plays a key role in maintaining the methylation state of these clusters. Moreover, only a limited number of genes showed MAC changes in the mutants with 19 losses and 7 gains in the *set1a*^−^/*set1b*^−^ mutant and 20 genes, 13 losses and 7 gains, in the *gcn5a*^−^/*gcn5b*^−^ mutant (**Supplementary Table 1**). These findings indicate that disruption of these activating histone modifications primarily alters individual methylation sites rather than the stability of entire MACs.

Finally, we assessed the transcriptional consequences of altered 6mA deposition in the *set1a*^−^/*set1b*^−^ and *gcn5a*^−^/*gcn5b*^−^mutants. Genes lacking 6mA, either at individual methylation sites or through the absence of MACs, exhibited the highest proportion of DEGs (Fisher’s exact test *p* < 0.01) (**Figure 4G and Supplementary Figure 4D**), supporting a role for 6mA in maintaining transcriptional stability. Furthermore, consistent with the association between 6mA and active transcription (42, 44, 51), genes that acquired de novo MACs were enriched among upregulated genes, whereas genes that lost MACs were preferentially associated with downregulation (Fisher’s exact test: Gain-Up *p* = 0.007 for *set1a*⁻/*set1b*⁻ and *p* = 0.25 for *gcn5a*⁻/*gcn5b,* Loss-Down *p* = 0.004 for *set1a*⁻/*set1b*⁻ and *p* = 0.0004 for *gcn5a*⁻/*gcn5b*) (**Figures 4H and 4I**). Overall, these results reveal a context-dependent interplay between histone modifications and 6mA deposition, whereby activating histone marks restrict ectopic adenine methylation, preserve MAC stability, and buffer transcriptional variation.

## DISCUSSION

Our study provides the first comprehensive mapping of activating histone modifications in a zygomycete fungus, revealing that H3K4me1, H3K4me3, and H3K27ac collectively define the euchromatic compartment of the *R. microsporus* genome. These marks delineate two functionally distinct euchromatic sub-compartments: one enriched for H3K4me3 at TSS and tightly co-localized with MACs, and another characterized by H3K4me1 and H3K27ac distributed across gene bodies and upstream regulatory regions. This bipartite organization mirrors the canonical distinction between active promoters marked by H3K4me3 and H3K27ac, and active enhancers or transcribed gene bodies enriched for H3K4me1 and H3K27ac, a regulatory logic first established in metazoans and later also found conserved in plants (52–54). The specific co-localization of H3K4me3, but not H3K4me1 or H3K27ac, with MACs suggests that the functional coupling between histone methylation and 6mA deposition is specific to the promoter-proximal chromatin and is likely tied to transcription initiation, consistent with the previously established association between 6mA and active gene expression in Mucorales (43).

The functional dissection of the Set1 and Gcn5 families reveals unexpected regulatory specialization among paralogs. *R. microsporus* encodes two Set1 paralogs: Set1a, responsible primarily for H3K4me3, and Set1b, which preferentially deposits H3K4me1, whereas related Mucorales such as *M. lusitanicus* encode a single Set1 ortholog (29). This functional specialization of Set1 paralog represents, to our knowledge, the first description of H3K4 methylation subfunctionalization in fungi and may reflect an evolutionary elaboration of chromatin regulatory capacity within the *Rhizopus* lineage. By contrast, Gcn5a and Gcn5b are functionally redundant for H3K27 acetylation; a significant reduction in H3K27ac was only observed in the *gcn5a*⁻/*gcn5b*⁻ double mutant. Such redundancy is commonly observed among essential chromatin regulators and likely reflects the central role of H3K27 acetylation in maintaining transcriptional programs across a broad range of genes (55).

The phenotypic consequences of disrupting these chromatin modifiers highlight their biological relevance in fungal development and pathogenicity. The *gcn5a*⁻/*gcn5b*⁻ double mutant exhibits severe growth defects and a complete loss of sporulation, while disruption of *set1a* and *set1b* reduces virulence and alters responses to cell wall and oxidative stress. These findings extend the well-established roles of Gcn5/SAGA in *Candida* species, where SAGA-mediated acetylation regulates antifungal resistance and virulence (56–58), to developmental processes in NDF. Furthermore, it is worth highlighting that, while disruption of genes encoding histone modifiers was feasible, attempts to disrupt any of the core components of the 6mA deposition machinery resulted in a lethal phenotype, consistent with their essential role (43). This observation suggests that 6mA plays a central and indispensable role in establishing and maintaining active chromatin domains, whereas histone modifiers may provide a more nuanced layer of transcriptional regulation for fine-tuning the modulation of transcriptional activity.

Our analyses also uncover an intricate interplay between histone modifications and DNA adenine methylation. In both the *set1* and *gcn5* mutants, we observed ectopic gains of 6mA at sites that are unmethylated in the wild-type strain, whereas global 6mA levels and symmetry remain largely conserved. This redistribution of 6mA, rather than a global elevation or decrease in methylation, suggests that the MTA1c complex responsible for 6mA deposition operates with a relatively fixed enzymatic capacity that is redirected to ectopic sites when specific histone marks are absent. A conceptually similar mechanism has been described in ciliates, where H3K4me3 limits the activity of MTA1c and restricts 6mA deposition (50). Our findings extend this principle to filamentous fungi and suggest that H3K4me3-dependent restriction of 6mA deposition may represent a conserved feature of epigenetic crosstalk across diverse eukaryotic lineages. One possible explanation is that H3K4me3-containing nucleosomes create a functional or steric barrier that limits inappropriate access of the 6mA methylation machinery to actively transcribed regions, thereby channeling methylation toward MACs at gene promoters.

The selective depletion of MACs at loci exhibiting high H3K27ac levels in the wild-type strain reveals an additional and mechanistically distinct role for lysine acetylation in the maintenance of established 6mA domains. While H3K4me3 appears to constrain the spread of 6mA to novel sites, H3K27ac seems to be required for the stability of a subset of pre-existing MACs. This observation is consistent with the well-established role of H3K27ac in promoting an accessible, nucleosome-depleted chromatin environment at regulatory regions (59), which may facilitate access to the MTA1c to symmetrically remethylate AT dinucleotides following DNA replication. Conversely, loss of H3K27ac may induce local chromatin compaction, thereby impairing the faithful maintenance of 6mA. The preferential loss of MACs at loci with high H3K27ac levels further supports a model in which 6mA maintenance at specific genomic regions depends on histone acetylation.

A direct consequence of 6mA redistribution is increased transcriptional variability. In both mutants, genes lacking 6mA in the wild-type strain show the greatest number of DEGs, whereas MAC-bearing genes remain comparatively stable, supporting a buffering role for 6mA in transcriptional regulation. The de novo acquisition of MACs in the *set1a^-^ /set1b^-^* mutant, predominantly associated with upregulation, may represent a compensatory epigenetic response to chromatin deregulation, whereby 6mA is redistributed to aberrantly activated loci to reinforce transcriptional stability. These findings suggest that the buffering function of 6mA arises through its coordinated interplay with histone modifications. The disproportionate vulnerability of short, densely methylated genes further indicates that protection by 6mA is finite. Although these genes display a high density of 6mA per base pair, their lower absolute number of methylated sites may be insufficient to preserve transcriptional stability when histone-mediated regulation is compromised.

Taken together, our findings support a multilayered epigenetic regulatory model in which active histone modifications and 6mA form an interdependent network ensuring transcriptional precision in *R. microsporus*. Symmetric 6mA plays a central role in establishing and maintaining active chromatin domains, whereas active histone modifications shape a chromatin landscape that permits accurate deposition and faithful maintenance of 6mA, thereby promoting stable transcriptional output. This positions *R. microsporus*, and NDF more broadly, as a tractable system for dissecting ancient epigenetic interactions that may have emerged early during eukaryotic evolution. Future studies aimed at elucidating the physical and functional interactions between the Set1 and Gcn5 complexes and the MTA1c will be necessary to determine whether this crosstalk is mediated by direct protein-protein interactions or indirectly through chromatin remodeling, and to establish whether the restriction of off-target 6mA deposition results from an active targeting mechanism or from the chromatin environment created by active histone modifications.

## METHODS

### Fungal growth conditions

The wild-type strain used in this study was *R. microsporus* ATCC 11559. Unless otherwise specified, strains were routinely grown on rich YPG medium (Yeast extract–Peptone–Glucose: 3 g/L yeast extract, 10 g/L peptone, 20 g/L glucose, 15 g/L agar) adjusted to pH 4.5, under continuous illumination at 30°C. Transformants derived from the auxotrophic strain UM33 (60) were selected using the *pyrF* or *leuA* selectable marker. *pyrF* transformants were grown in Minimal Media with Casamino acids (MMC), whereas *leuA* transformants were selected on Yeast Nitrogen Base media (YNB) (61). Following electroporation, protoplasts were recovered in ice-cold YPG supplemented with 0.5 M Sorbitol for 90 min, collected by centrifugation at 80 × g, and resuspended in YNB containing 0.5 M sorbitol. Protoplasts were then plated onto MMC or YNB supplemented with 0.5 M sorbitol and adjusted to pH 3.2. Plates were incubated at 30 °C for 4-5 days until transformant colonies were visible. To obtain homokaryotic transformants, selected transformants were serially passaged several times on the corresponding selective medium (MMC or YNB, pH 3.2) depending on the selectable marker used. MMC and YNB media were supplemented with thiamine and niacin as previously described (62, 63).

### Mutant strain generation

*R. microsporus* mutant strains were generated via CRISPR-Cas9–mediated gene disruption using microhomology-directed repair DNA templates (62, 63). For the generation of single *set1 and gcn5* mutants, repair templates containing the *pyrF* selectable marker flanked by 38-nt microhomology arms were used to disrupt the targeted loci. Homology-directed repair was promoted by Cas9-induced double-strand breaks (DSB) in the uracil and leucine auxotrophic strain UM33 (60). Target-specific CRISPR crRNAs (Alt-R^TM^ CRISPR-Cas9 crRNA, IDT) were designed for each gene (Supplementary Table 2) and annealed to tracRNA (Alt-R^TM^ CRISPR-Cas9 tracRNA, IDT) to generate the guide RNAs (gRNAs). The resulting gRNAs were assembled in vitro with recombinant Cas9 nuclease (Alt-R^TM^ S.p. Cas9 nuclease, IDT) to form ribonucleoprotein complexes prior to transformation. Gene disruptions and homokaryotic status of the resulting transformants were validated by PCR using primer pairs flanking the targeted integration site, allowing amplification of both WT and disrupted alleles when present. All primers used for repair-template construction and PCR validation are listed in the **Supplementary Table 3**. Genomic DNA was extracted by phenol/chloroform extraction (64), and all PCR amplifications were performed with Herculase II Fusion DNA polymerase (Agilent), optimizing annealing temperatures to each primer pair following the manufacturer’s recommendations.

### ChIP-seq

*R. microsporus* ChIP-seq experiments were conducted as previously described (43). Briefly, mycelium was crosslinked with formaldehyde (1%) and quenched with glycine at a final concentration of 135 mM for 10 min. After flash-freezing with liquid nitrogen, 180 mg of frozen mycelium per ChIP experiment were ground to a fine powder, which was resuspended in 900 µL ChIP Lysis buffer (50 mM HEPES pH 7.4, 150 mM NaCl, 1% Tritton X-100, 0.1% DOC, and 1mM EDTA) and 10 µL of protease inhibitor cocktail. The lysates were sonicated using a Bioruptor Pico sonicator (Diogenode) with 30 s ON/OFF pulses for 25 cycles in a thermostated bath (4 °C). After sonication, samples were centrifuged at 12.000 × g for 8 min at 4 °C and transferred and divided into 3 tubes: 100 µL were frozen at -20 °C for input DNA control, 400 µL were used for antibody immunoprecipitation (IP) and 400 µL were used for beads only control. Beads were equilibrated with dilution buffer (50 mM HEPES pH 7.4, 150 mM NaCl, and 1 mM EDTA). For H3K4me1, H3K4me3, H3K27ac, and RNA Pol II experiments, 50 µL of equilibrated Dynabeads protein A magnetic beads (ThermoFisher) were added to mock beads only samples and another 50 µL were conjugated with the selected ChIP-grade antibodies (Anti-H3K4me1: Abcam # ab8895; anti-H3K4me3: Abcam #ab213224; anti-H3K27ac: Abcam #ab4729; and anti-RNA Pol II: Active Motif #ab_2732926) at 4 °C for 2 h. After washing again with dilution buffer, antibody-conjugated Dynabeads were added to the IP samples. Both IP and bead control were incubated overnight at 4 °C. After incubation, the beads were washed twice with 1 mL of Low Salt Wash Buffer (20 mM Tris pH 8.0, 150 mM NaCl, 0.1% SDS, 1% Triton X-100, and 2 mM EDTA), twice with 1 mL of High Salt Wash Buffer (20 mM Tris pH 8.0, 500 mM NaCl, 0.1% SDS, 1% Triton X-100, and 2 mM EDTA), once with LiCl Wash Buffer (10 mM Tris pH 8.0, 1 mM EDTA, 0.25 M LiCl, 1% NP40, and 1% DOC), and finally once with TE (10mM Tris pH 8.0 and 1 mM EDTA). DNA was eluted from beads by adding 250 µL of TES Buffer (50 mM Tris pH 8.0, 10 mM EDTA, 1% SDS) and incubating for 10 min at 65 °C (twice). The final 500 µL eluate and the input DNA samples were submitted to reverse crosslinking by incubating overnight at 65 °C in the presence of 0.2 M NaCl. Samples were treated with Proteinase K (Merck) at a final concentration of 1 mg/mL and RNAse A (Merck) at a final concentration of 1 mg/mL for 3 h at 65 °C. DNA was purified by adding 1/2 volume of phenol and 1/2 volume of chloroform:isoamyl alcohol (24:1). After centrifugation (2 min at 16000 × g), the aqueous phase was transferred to a new tube and 1 volume of chloroform:isoamyl alcohol (24:1) was added. After centrifugation (2 min at 16000 × g), the aqueous phase was transferred to a new tube and 1/10 volume of 3 M sodium acetate, 1 volume of ethanol (96%), and 20 µg of glycogen (ThermoFisher) were added. The tubes were incubated at -20 °C overnight and DNA was precipitated by centrifugation at 4 °C, 10 min at 16000 × g. DNA was washed twice with 70% ethanol and dried at room temperature for 10 min. DNA was resuspended with 30 µL of ddH_2_0. Purity and DNA quantity were analyzed with Nanodrop 2000 (Thermo Scientific) and Qubit (Thermo Scientific). Purified DNA samples were used to construct libraries and to be sequenced (Illumina NovaSeq 6000, 150-bp paired-end reads) by Novogene. Sequencing adapters were trimmed using Cutadapt (v4.0) (65). Trimmed reads were mapped to the *R. microsporus* v2.0 reference genome (available at https://mycocosm.jgi.doe.gov/Rhimi59_2/Rhimi59_2.home.html) using bwa-mem for medium and long reads with default parameters (v0.7.17) (66). BAM files were sorted and deepTools bamCoverage function (67) (bin size = 10, CPM normalization) was used to visualize and manually inspect tracks on IGV (v2.12.13) (68). bamCompare function (bin size = 10, CPM normalization) was used to compute ratios between IP and input samples. The resulting bigwig or bedgraph files were used for downstream analysis. To assess the specificity of the immunoprecipitations, the mock beads-only signal was subtracted, and the resulting tracks were manually analyzed on IGV (v2.12.13) (68). MACS2 software (v2.2.7.1) (69) was used to call peaks for each ChIP-seq experiment (effective genome size = 27394811, FDR cut-off = 0.05) on narrow peaks mode for H3K4me1, H3K4me3, H3K27ac, and RNA Pol II.

### RNA-seq

Total RNA was extracted using the NZY Total RNA Isolation Kit (NZYTech) following the manufacturer’s instructions and treated with DNase I (Sigma, On-Column DNase I Treatment Set). RNA integrity was assessed using a Bioanalyzer 2100 system (Agilent Technologies). Sequencing libraries were prepared and sequenced by Novogene. Raw sequencing read were processed using Trimmomatic v0.39 (70) to remove adapter sequences, low-quality bases, short reads (< 20 nt), and sequencing artifacts. Filtered reads were aligned to the *R. microsporus* v2.0 reference genome using STAR v2.7.11a (71). Individual read count matrices were generated from the resulting Binary Alignment Map (BAM) files using featureCounts v2.0.8 (72) and further processed for differential expression analysis between mutant and wild-type strain with DESeq2 v2.11.40.8 (73). Genes were considered differentially expressed when they displayed an adjusted *p* < 0.05 and an absolute log₂ fold change (|log₂FC|) ≥ 2.

### PacBio sequencing and 6mA analysis

6mA base modifications were detected with the PacBio SMRT analysis platform (cromwell.workflows.pb_basemods). 10 µg of genomic DNA was sheared to >10 kb using Covaris g-Tubes. The sheared DNA was treated with exonuclease to remove single-stranded ends and DNA damage repair mix, followed by end repair and ligation of blunt adapters using SMRTbell Template Prep Kit 1.0 (Pacific Biosciences). The library was purified with AMPure PB beads and size-selected with BluePippin (Sage Science) at >10 kb cutoff size. PacBio Sequencing primer was then annealed to the SMRTbell template library and sequencing polymerase was bound to them using Sequel Binding kit 3.0. The prepared SMRTbell template libraries were then sequenced on a Pacific Biosciences Sequel sequencer using v3 sequencing primer, 1 M v3 SMRT cells, and Version 3 sequencing chemistry with 1 × 600 sequencing movie run times. Raw reads were filtered using SFilter, to remove short reads and reads derived from sequencing adapters. Filtered reads were aligned to the reference genome of *R. microsporus* v2 (https://mycocosm.jgi.doe.gov/Rhimi59_2/Rhimi59_2.home.html), using BLASR (1.5.0)122. Modified sites were then identified through kinetic analysis of the aligned DNA sequence data123. Detected 6mA sites were further filtered with a minimum 15× per-strand coverage to remove potential false positives and select only 6mA sites with >25 mQv. Methylated adenine sites were called using the SMRT Link pipeline and converted to binary BigWig format (value = 1 per site, 1-bp resolution) using bedGraphToBigWig (UCSC). MACs were defined as previously described (42, 44). Gene-level 6mA density was computed as the number of methylated sites per kilobase of gene body, and dynamic changes between WT and mutant strains were classified as Gained or Lost when density changed ≥ 2-fold, or as Stably low when both WT and mutant had ≤ 1 site. Average 6mA profiles over genomic regions were computed with deepTools (67) and statistical comparisons were performed with the Wilcoxon signed-rank test.

### Protein extraction and western blot analysis

For protein extracts, 100 mg of mycelium were lysed in 300 µL of breaking buffer (8 M urea, 5% w/v SDS, 40 mM Tris–HCl pH 6.8, 0.1 mM EDTA, 0.4 mg/mL bromophenol blue, 1% β-mercaptoethanol, and 5 mM PMSF) using 0.5 mm silica beads in a FastPrep-24™ homogenizer (three cycles of 30 s at 4 m/s, with cooling on ice between cycles). Cell debris was removed by centrifugation at 14,000 rpm for 10 min at 4 °C, and the supernatant was transferred to a clean tube on ice. The pellet was resuspended in 50 µL of breaking buffer, vortexed for 1 min, and centrifuged again at 14,000 rpm for 5 min at 4 °C. The resulting supernatant was pooled with the previous one.

Protein concentration was determined using a Qubit Fluorometer (Invitrogen, Thermo Fisher Scientific), and 150 µg of total protein was separated on NuPAGE™ 4–12% Bis-Tris gradient gels (Invitrogen) using MOPS SDS NuPAGE™ running buffer (Invitrogen) at a constant voltage of 200 V for 45 min. Proteins were transferred onto nitrocellulose membranes using a semi-dry electroblotting System (Sigma-Aldrich). Membranes were probed with antibodies against H3 (Abcam, ab1791), H3K4me1 (Abcam, ab8895), H3K4me2 (Abcam, ab176878), H3K4me3 (Abcam, ab8580), and H3K27ac (Abcam, ab4729). Immunoreactive signals were quantified by densitometric analysis using ImageJ (74). Quantification was performed from three different replicates and data are presented as the mean ± SD.

### Phenotypic characterization

For sporulation and growth assays, 100 spores of each strain were inoculated at the center of YPG agar plates (pH 4.5) and incubated at 26 °C under illuminated conditions for 96 h. Sporulation was determined by quantifying spores produced using a Neubauer chamber and normalizing to colony size. Colony diameter was recorded every 24 h for four days. For stress susceptibility assays, 100 fresh spores were inoculated in the center of YPG agar plates (pH 4.5) supplemented with SDS (0.001-0.002%) or H_2_O_2_ (1-2 mM). Plates were incubated at 26°C for 72 h, and the diameter of the colony was registered. Growth inhibition (%) was calculated as [(control diameter – treated diameter) / (control diameter)] x 100. For virulence assays in *G. mellonella,* groups of 20 adult larvae were injected through the last prolegs with 10^6^ spores in 20 μL of insect physiological saline solution (IPS), as previously described (75). Larvae injected with IPS were employed as a control group. Following infection, larvae were incubated at 30 °C, and survival was recorded every 24 h for six 6 days. Average survival data were obtained from three independent experiments and were analyzed using the log-rank (Mantel-Cox) test with GraphPad Prism 10 software. Data with *p* < 0.05 were considered statistically significant.

### Statistical analysis

Statistical details are detailed in the Results, Figures, and Figure legends, including the number of biological and technical replicates as well as the dispersion and precision measures (mean and SD). Statistical analyses were performed using GraphPad Prism 9 (https://www.graphpad.com). Data normality was analyzed using the Shapiro-Wilk normality test with a significance level (alpha) of 0.05. Densitometry comparisons were performed using Welch’s t-test *p* threshold of < 0.05. Wilcoxon rank test was conducted for each bin and the P-value corrected for multiple comparisons was calculated using deepStats package (https://github.com/gtrichard/deepStats) (v0.3.31). TPMs differences in different sets of genes were evaluated using the Kolmogorov-Smirnov test, with a *p* < 0.05 as the cutoff to determine statistically significant differences. Experimental data on colony growth and sensitivity were generated from two biological replicates and two technical replicates for each experiment. Each graph represents the mean ± SD from assay results. Data were analyzed to determine the statistical significance by One-way or Two-way ANOVA with Tukey’s post hoc test (*p* < 0.05). The significant differences between each data group are indicated in each graph with significant markers (** *p* < 0.001, **** *p* < 0.0001, ns; not significant. Virulence assay results were analyzed using the log-rank (Mantel-Cox) test with GraphPad Prism 9 software. Data with p ≤ 0.05 were considered statistically significant.

## ACKNOWLEDGMENTS

This research was supported by the grant PID2024-160088NB-I00 to F.E.N. and V.G, funded by MICIU/AEI/10.13039/501100011033 and by ERDF/EU. The work (10.46936/10.25585/60001127) was conducted by the US Department of Energy Joint Genome Institute (https://ror.org/04xm1d337), a DOE Office of Science User Facility, supported by the Office of Science of the US Department of Energy under Contract No. DE-AC02-05CH11231.

## DATA AVAILABILITY

All data used in this study have been deposited in public repositories. RNA-seq and ChIP-seq data have been deposited in GEO under the accession numbers GSE326211, GSE326209, GSM8509639, and GSM8509642. PacBio sequencing data have been deposited at the Sequence Read Archive (SRA) under the accession numbers SRP701151, SRP701152, and SRP701153.

## CODE AVAILABILITY

Custom scripts generated for data processing are available at the GitHub repository (https://doi.org/10.5281/zenodo.19219904).

## AUTHOR CONTRIBUTIONS

C.L. supervised experiments, processed the data and conducted analyses, prepared the figures and tables, designed and coordinated the project, and wrote the manuscript with significant input from F.E.N. and V.G. M.O.C. designed and coordinated the project, generated knockout mutant strains, conducted phenotypic characterization, and participated in figure preparation. N.N.M. conducted Western blot experiments and generated samples for sequencing. G.T. developed scripts and code for data processing, analysis and visualization, prepared figures and tables and participated in manuscript writing. S.J.M. participated in 6mA data processing. V.N. managed the project. E.N. managed the project and provided materials. I.V.G. supervised and coordinated the project. V.M.C supervised the investigation and provided resources. F.E.N. analyzed the results and supervised the study. V.G. analyzed the results and designed, supervised, and coordinated the project.

## SUPPLEMENTARY FIGURE AND TABLE LEGENDS

**Supplementary Figure 1.** (**A**) Genomic tracks of genes (light brown), repeats (purple-blue), 6mA (pink), H3K4me1 (light blue), H3K4me3 (dark blue), H3K27ac (green), H3K9me3 (dark purple), RNA Pol II (orange), and RNA-seq (yellow) across scaffolds 1-28. Scale = 500 kb. (**B**) RNA Pol II occupancy over genes. Genes were classified into four quartiles (Q1-Q4) based on their expression levels in the wild-type strain. Genes were sized to 2 kb and occupancy was also computed 1.5 kb upstream and downstream the gene body. (**C**) H3K4me1 (top), H3K4me3 (mid) and H3K27ac (bottom) occupancy over quartile-classified genes according to their expression levels. Genes were sized to 2 kb and occupancy was also computed 1.5 kb upstream and downstream the gene body.

**Supplementary Figure 2.** (**A**) Top, schematic representation of *set1a* and *set1b* gene disruption performed using CRISPR/Cas9. Arrows indicate the primers used for mutant confirmation. Bottom, agarose gel electrophoresis of PCR products confirming correct gene disruption and homokaryosis status of the generated mutants. (**B**) Left, schematic representation of *gcn5* gene disruption performed using CRISPR/Cas9. Arrows indicate the primers used for mutant confirmation. Right, gel electrophoresis of PCR products confirming the mutation and homokaryosis status of the generated mutants. (**C**) and (**F**) Images of *R. microsporus* wild-type (WT) strain and *set1* and *gcn5* mutants growing at different SDS and H_2_O_2_ concentrations. (**D**), (**E**), (**G**), and (**H**) Growth inhibition of the WT strain and indicated *set1* and *gcn5* mutants at different SDS and H_2_O_2_ concentrations, expressed as the percentage of growth inhibition relative to the untreated control after 72 h of incubation. Data represent the mean ± SD of two independent biological replicates, each including two technical replicates. Two-way ANOVA was used for statistical analysis, with Tukey multiple comparison test (*p* < 0.05). Statistically significant differences obtained by ANOVA are shown in each graph as follows: \*\**p* < 0.01; **** *p* < 0.0001; ns, not significant.

**Supplementary Figure 3. (A)** Genomic tracks of genes (light brown), repeats (purple-blue), 6mA (pink), H3K4me1 (light blue), H3K4me3 (dark blue), H3K27ac (green), and H3K9me3 (dark purple) for wild-type and *set1* and *gcn5* double mutant strains across scaffolds 1-28. Scale = 500 kb. (**B**) RNA Pol II occupancy over *R. microsporus* genes. Profiles are shown for WT strain (black), *set1a*^-^/*set1b*^-^ mutant (blue), and *gcn5*a^-^/*gcn5b^-^*mutant (green). Genes were sized to 2 kb and occupancy was also computed 1.5 kb upstream and downstream the gene body.

**Supplementary Figure 4. (A)** Boxplots comparing gene length, number of absolute methylated sites, and 6mA density among change categories (Gained, Lost, Unchanged, Stably low) for *set1a*⁻/*set1b*⁻ (top) and *gcn5a*⁻/*gcn5b*⁻ (bottom) mutants. Welch’s t-test was used for pairwise comparisons; significance levels are indicated (* *p* < 0.05, *** *p* < 0.01). (**B**) Average ChIP-seq signal profiles (ratio to input) for H3K4me1, H3K4me3 and H3K27ac in WT, *set1a*⁻/*set1b*⁻ and *gcn5a*⁻/*gcn5b*⁻ mutants over MACs that were lost in each mutant. Solid line = WT, dashed line = mutant. Each region was extended to ±1.5 kb around the MAC body. (**C**) Violin plots showing mean WT H3K27ac signal (ChIP-seq ratio to input) for all MACs (n = 7,068) versus MACs lost in *gcn5a*⁻/*gcn5b*⁻ (n = 258). Wilcoxon rank-sum test; *** *p* < 0.01. (**D**) Upregulated (red) and downregulated (blue) DEGs (|log₂FC| ≥ 2, FDR < 0.05) stratified by MAC status in each mutant. Fisher’s exact test no MAC vs MAC genes *p* = 2.4 × 10^-298^ for *set1a*⁻/*set1b*⁻ and *p* = 1.2 × 10^-207^ for *gcn5a*⁻/*gcn5b*⁻.

**Supplementary Table 1.** DEGs with MAC gains and losses in the *set1a*⁻/*set1b*⁻ and *gcn5a*⁻/*gcn5b*⁻ mutants.

**Supplementary Table 2.** crRNAs used in this work.

**Supplementary Table 3.** Primers used in this work.

**Supplementary Table 4.** Fungal strains used in this work

**Supplementary File 1**. Spreadsheets containing differentially expressed genes and MACs localization and methylation state for each gene in WT and mutant strains.

