## Supplementary information for "Active histone modifications fine-tune DNA N-6 methyladenine deposition and maintain transcriptional stability"

Supplementary Figure 1

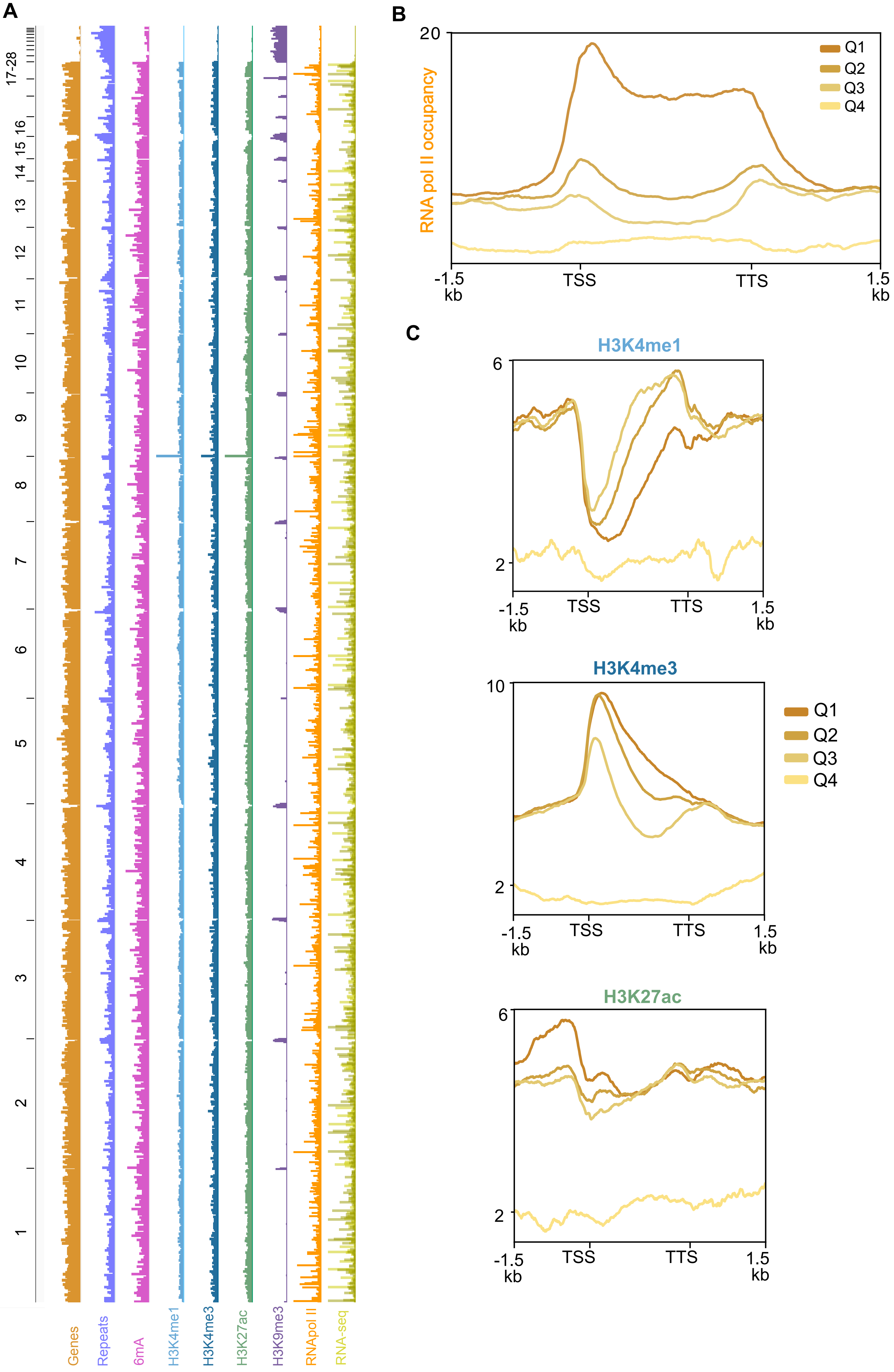

Supplementary Figure 2

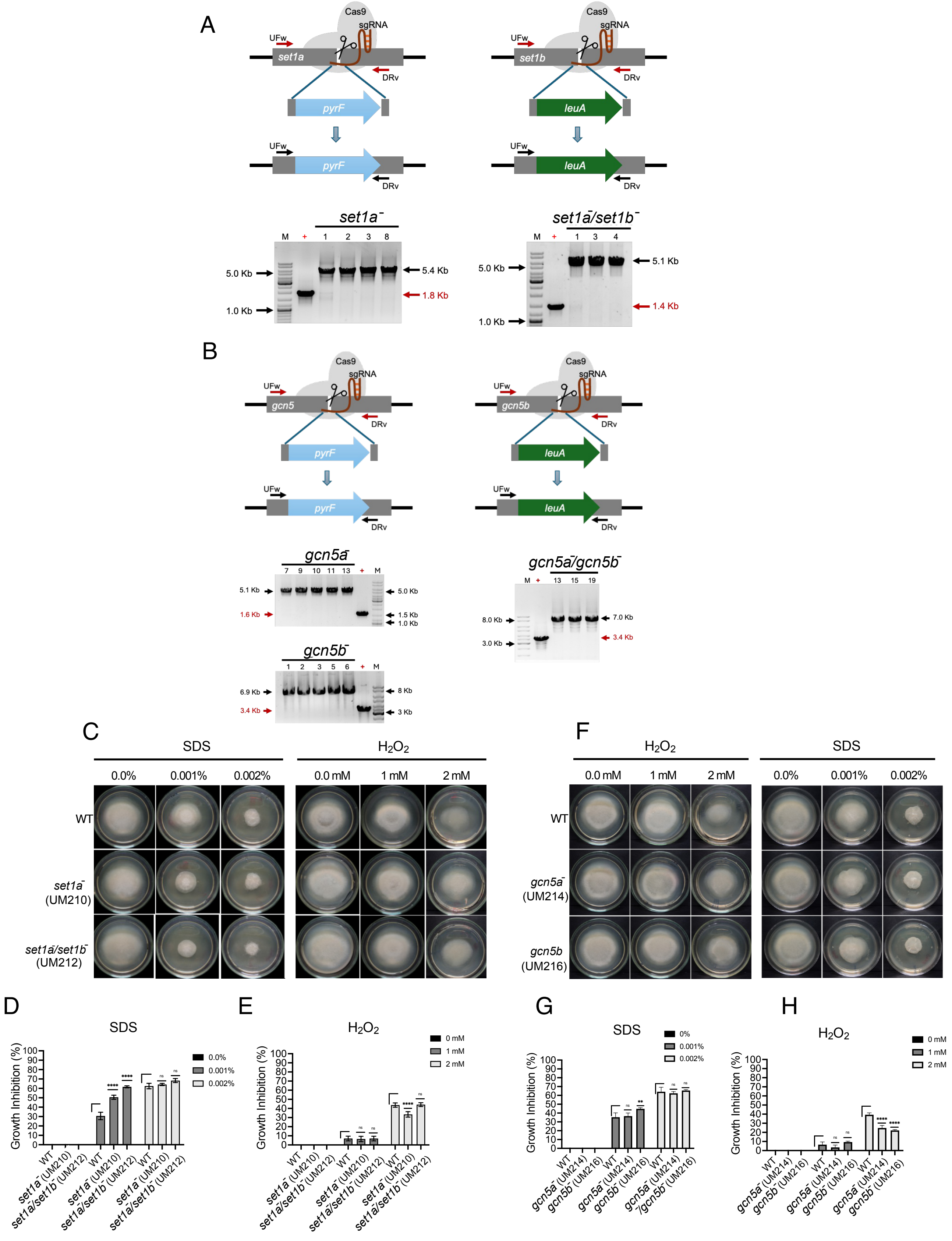

Supplementary Figure 3

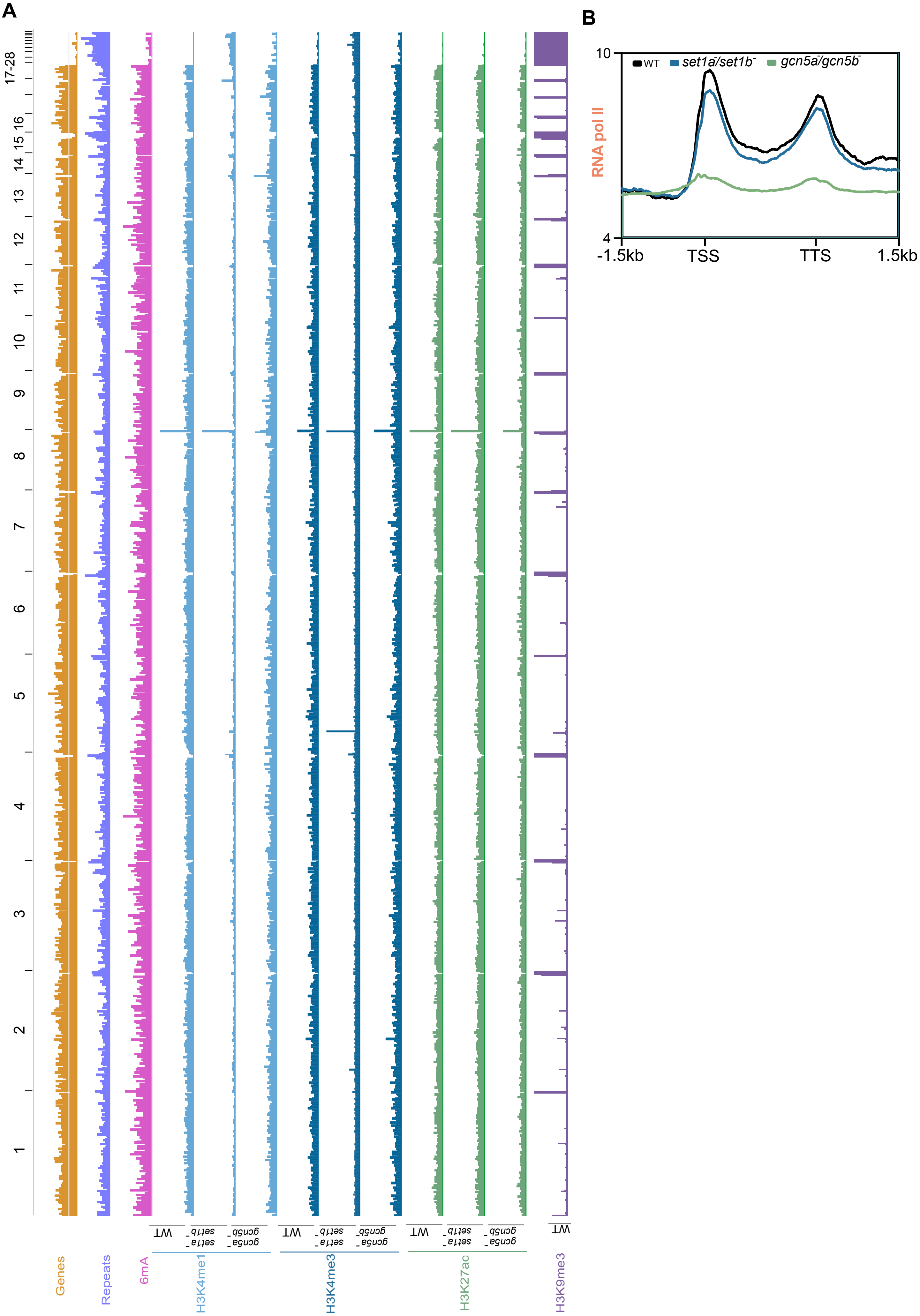

Supplementary Figure 4

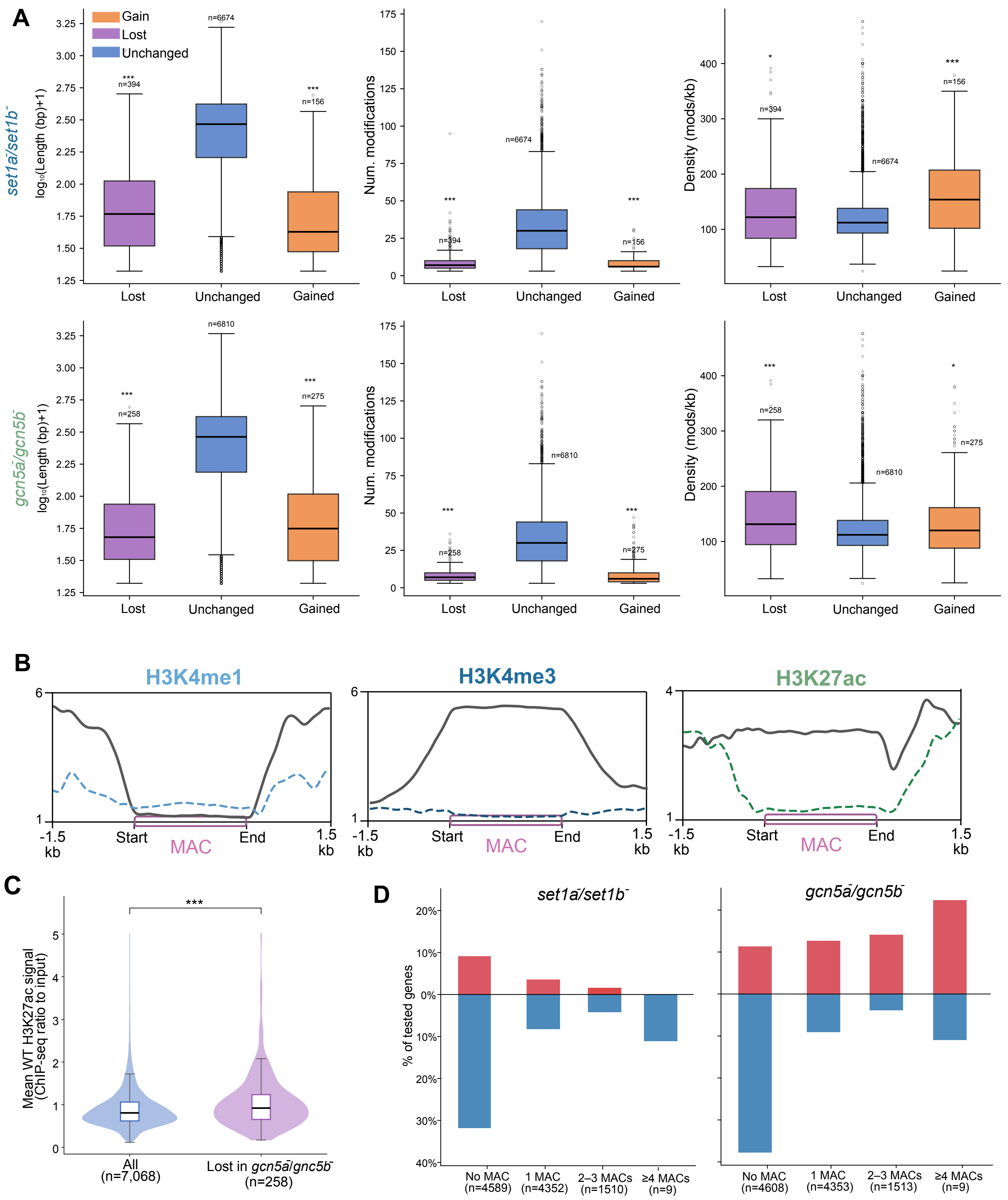

**Table S1**. MAC gains and losses in Set1 and Gcn5 mutants.

| geneID | MAC WT | MAC Set1 Mut | MAC Gcn5 Mut | Category set1 | Category gcn5 |
|---|---|---|---|---|---|
| **1883390** | 1 | 3 | 1 | Gained | Unchanged |
| **1891719** | 1 | 3 | 1 | Gained | Unchanged |
| **1898002** | 1 | 3 | 1 | Gained | Unchanged |
| **1926216** | 0 | 2 | 1 | Gained | Unchanged |
| **1903462** | 1 | 3 | 2 | Gained | Unchanged |
| **1909116** | 1 | 3 | 2 | Gained | Unchanged |
| **1915902** | 1 | 3 | 2 | Gained | Unchanged |
| **1806500** | 3 | 1 | 1 | Lost | Lost |
| **1808971** | 2 | 0 | 0 | Lost | Lost |
| **1859702** | 2 | 0 | 0 | Lost | Lost |
| **1866803** | 3 | 1 | 1 | Lost | Lost |
| **1891963** | 2 | 0 | 0 | Lost | Lost |
| **1904865** | 2 | 0 | 0 | Lost | Lost |
| **1910024** | 3 | 1 | 1 | Lost | Lost |
| **1926023** | 2 | 0 | 0 | Lost | Lost |
| **1983927** | 5 | 2 | 2 | Lost | Lost |
| **1426148** | 3 | 1 | 2 | Lost | Unchanged |
| **1806378** | 3 | 1 | 3 | Lost | Unchanged |
| **1813390** | 3 | 1 | 2 | Lost | Unchanged |
| **1838378** | 3 | 1 | 2 | Lost | Unchanged |
| **1871241** | 3 | 1 | 3 | Lost | Unchanged |
| **1878496** | 2 | 0 | 1 | Lost | Unchanged |
| **1879882** | 2 | 0 | 2 | Lost | Unchanged |
| **1914785** | 2 | 0 | 1 | Lost | Unchanged |
| **1915346** | 2 | 0 | 1 | Lost | Unchanged |
| **1976232** | 3 | 1 | 3 | Lost | Unchanged |
| **1881745** | 1 | 1 | 3 | Unchanged | Gained |
| **1902972** | 0 | 0 | 2 | Unchanged | Gained |
| **1950293** | 0 | 0 | 2 | Unchanged | Gained |
| **1953907** | 0 | 1 | 2 | Unchanged | Gained |
| **1917787** | 1 | 2 | 3 | Unchanged | Gained |
| **1960611** | 1 | 2 | 3 | Unchanged | Gained |
| **1976649** | 1 | 2 | 3 | Unchanged | Gained |
| **1887470** | 3 | 2 | 1 | Unchanged | Lost |
| **1906985** | 2 | 1 | 0 | Unchanged | Lost |
| **1920385** | 4 | 3 | 1 | Unchanged | Lost |
| **1966863** | 2 | 1 | 0 | Unchanged | Lost |

**Table S2**. crRNA sequences used in this study.

| **crRNA name** | **Sequence 5' ⟶ 3'** |
|---|---|
| set1a 1 (set1a_sRNA1) | ATTCGAGACACCACTATTAC AGG |
| set1b 2 (set1b_sRNA2) | GATGATCCTAGTGATGCCTT GGG |
| gcn5a 3 (gcn5a_sRNA3) | GTACAGGTCGTGCTCAATGA TGG |
| gcn5b 4 (gcn5b_sRNA4) | GAAGTGGGTGGGGTATATCA AGG |

**Table S3**. Oligonucleotide sequences used in this study.

| **Primer name** | **Sequence 5'⟶3'** |
|---|---|
| set1a 1-UFw | CAAACTGCTGCCGACAAGAC |
| set1a 1-DRv | CATGTTGAGGTGAAGGATACAGA |
| set1b 2-UFw | ACTGGCATATCCTATTCCCTTC |
| set1b 2-DRv | CAAGGTATCTTGTCCGCTTCA |
| gcn5a 3-UFw | ACCCTGATATTACAAAGCACAAGG |
| gcn5a 3-DRv | CTAAACGATCAACGGGTATAATCT |
| gcn5b 4-UFw | GCGCTAGCGTCTTTGTTGTA |
| gcn5b 4-DRv | CGTCTCGCTCTTCTGCTTTT |
| Templ_set1a 1_F | GCTATATCGATGATTATAGCTATTCGAGACACCACTATTCCT CCATAAGAATTTGACAG |
| Templ_set1a 1_R | CTAGATGAATAATACTGAGGTGGAGACCTGCTCCTGTATGA TAAAACGAAGATGTGGCTGTC |
| Templ_set1b 2_F | TTGGATTCATTAAAAAAGGAAGATGATCCTAGTGATGCTCCT CCATAAGAATTTGACAG |
| Templ_set1b 2_R | TAGGGCTGTATGCTCTTGTTCTAGCTGATTGACCCAAGTGAT AAAACGAAGATGTGGCTGTC |
| Templ_gcn5a 3_F | AAGGGAAGGAAGCATTCGAGTACAGGTCGTGCTCAATGTC CTCCATAAGAATTTGACAG |
| Templ_gcn5a 3_R | TAGGCCTGTCAAGAGGATCATACTTTCGCGCGCTCCATTGA TAAAACGAAGATGTGGCTGTC |
| Templ_gcn5b 3_F | AGAGATCACACTCGACAAGAGGAAGTGGGTGGGGTATATCC TCCATAAGAATTTGACAG |
| Templ_gcn5b 3_R | TCTCACCTGCATAATCGTTCCTCCTTCATAATCCTTGATGATA AAACGAAGATGTGGCTGTC |

**Table S4**. *R. microsporus* strains employed in this work.

| **Strain name** | **Genotype** | **Source** |
|---|---|---|
| ATCC 11559 | WT | ATCC |
| UM33 | *pyrF-*, *leuA*$^{-}$ | *Tahiri et al*. 2026 |
| UM210 | *set1a::pyrF*$^{+}$, *leuA*$^{-}$ | This work |
| UM211 | *set1a::pyrF*$^{+}$, *leuA*$^{-}$ | This work |
| UM212 | *set1a::pyrF*$^{+}$, *Δset1b::leuA*$^{+}$ | This work |
| UM213 | *set1a::pyrF*$^{+}$, *Δset1b:: leuA*$^{+}$ | This work |
| UM214 | *gcn5a:: pyrF*$^{+}$, *leuA*$^{-}$ | This work |
| UM215 | *gcn5a:: pyrF*$^{+}$, *leuA*$^{-}$ | This work |
| UM216 | *gcn5b:: pyrF*$^{+}$, *leuA*$^{-}$ | This work |
| UM217 | *gcn5b:: pyrF*$^{+}$, *leuA*$^{-}$ | This work |
| UM218 | *gcn5a::pyrF*$^{+}$, *Δgcn5b:: leuA*$^{+}$ | This work |
| UM219 | *gcn5a::pyrF*$^{+}$, *Δgcn5b:: leuA*$^{+}$ | This work |
